# Parallel and divergent morphological adaptations underlying the evolution of jumping ability in ants

**DOI:** 10.1101/2023.03.11.531676

**Authors:** Lazzat Aibekova, Roberto A. Keller, Julian Katzke, Daniel M Allman, Francisco Hita Garcia, David Labonte, Ajay Narendra, Evan P. Economo

## Abstract

Jumping is a rapid locomotory mode widespread in terrestrial organisms. However, it is a rare specialization in ants. Forward jumping has been reported within four distantly related ant genera: *Gigantiops*, *Harpegnathos*, *Myrmecia*, and *Odontomachus*. The temporal engagement of legs/body parts during jump, however, varies across these genera. It is unknown what morphological adaptations underlie such behaviors, and whether jumping in ants is solely driven directly by muscle contraction or additionally relies on elastic recoil mechanism. We investigate the morphological adaptations for jumping behavior by comparing differences in the locomotory musculature between jumping and non-jumping relatives using x-ray micro- CT and 3D morphometrics. We found that the size-specific volumes of the trochanter depressor muscle (*scm6*) of the middle and hind legs are 3-5 times larger in jumping ants, and that one coxal remotor muscle (*scm2*) is reduced in volume in the middle and/or hind legs. Notably, the enlargement in the volume of other muscle groups is directly linked to the legs or body parts engaged during the jump. Furthermore, a direct comparison of the muscle architecture revealed two significant differences between in jumping versus non-jumping ants: First, the relative Physiological Cross-Sectional Area (PCSA) of the trochanter depressor muscles of all three legs were larger in jumping ants, except in the front legs of *O. rixosus* and *M. nigrocincta*; second, the relative muscle fiber length was shorter in jumping ants compared to non-jumping counterparts, except in the front legs of *O. rixosus* and *M. nigrocincta*. This suggests that the difference in relative muscle volume in jumping ants is largely invested in the area (PCSA), and not in fiber length. There was no clear difference in the pennation angle between jumping and non-jumping ants. However, the length of hind legs relative to body length was longer in jumping ants. Based on direct comparison of the observed vs. possible work and power output during jumps, we surmise that direct muscle contractions suffice to explain jumping performance, in two species, but elastic recoil is likely important in one. We suggest that increased investment in jumping-relevant musculature is a primary morphological adaptation that separates jumping from non-jumping ants. These results elucidate the common and idiosyncratic morphological changes underlying this rare adaptation in ants.

## INTRODUCTION

Understanding the coevolution between morphology and behavior is one of the central challenges in evolutionary biology. Changes in environment and competition for resources can trigger new innovations in behavior, which in turn lead to morphological and physiological adaptations (Wcislo, 1989). Similar desired functionality can either result in convergent evolution of morphology, especially when a limited range of forms is readily accessible to evolution (McGhee, 2011), or manifest itself in many-to-one mapping (Moen, 2019; Wainwright et al., 2005), where the same functional trait is achieved by a diversity of morphological “design solutions”.

The evolution of jumping behaviors provides an excellent opportunity to study the relationship between function and morphology, because it often involves a combination of structural transformations, e.g., enlargement of muscular volumes and associated tendons, expansion of skeletal elements for larger attachment areas, and adaptations for elastic energy storage (Gorb, 2004; Ogawa & Yoshizawa, 2017). Furthermore, previous studies on insects have revealed a variety of distinct morphological designs that promote jumping. For example, locusts (Bennet Clark, 1975), click beetles (Bolmin et al., 2021) and froghoppers (Burrows, 2006) use catapult mechanisms to jump; other insects, such as mantises (Sutton et al., 2016), bush crickets (Burrows & Morris, 2003) and moths (Burrows & Dorosenko, 2015), in turn, rely on direct muscle actuation without additional contributions from elastic elements.

Despite the astounding diversity of ants, the ability to jump is rare; it has been reported in only six genera (Ali et al., 1992; Baroni Urbani et al., 1994; Sorger, 2015; Tautz et al., 1994; Wheeler, 1922). Jumping ants can be divided into two broad groups: prosalient, that is forward jumping using legs; and retrosalient, i.e., backward jumping using mandibles (Wheeler, 1922). Retrosalience is observed in trap-jaw ants such as *Odontomachus* Latrielle, 1804, *Strumigenys* Smith, F., 1860 and *Anochetus* Mayr, 1861, which use large muscles in their head to store elastic energy in both tendons and the head capsule (Sutton et al., 2022), which is then rapidly released, resulting in an impact between mandibles and the ground, and upward propulsion. Prosalient ants, on the other hand, use their legs to power a directed forward jump. Prosalience has evolved in four distantly related ant genera (see Fig 1.): *Harpegnathos*, *Gigantiops*, *Myrmecia,* and *Odontomachus* (Ali et al., 1992; Baroni Urbani et al., 1994; Sorger, 2015; Tautz et al., 1994).

**Figure 1.**
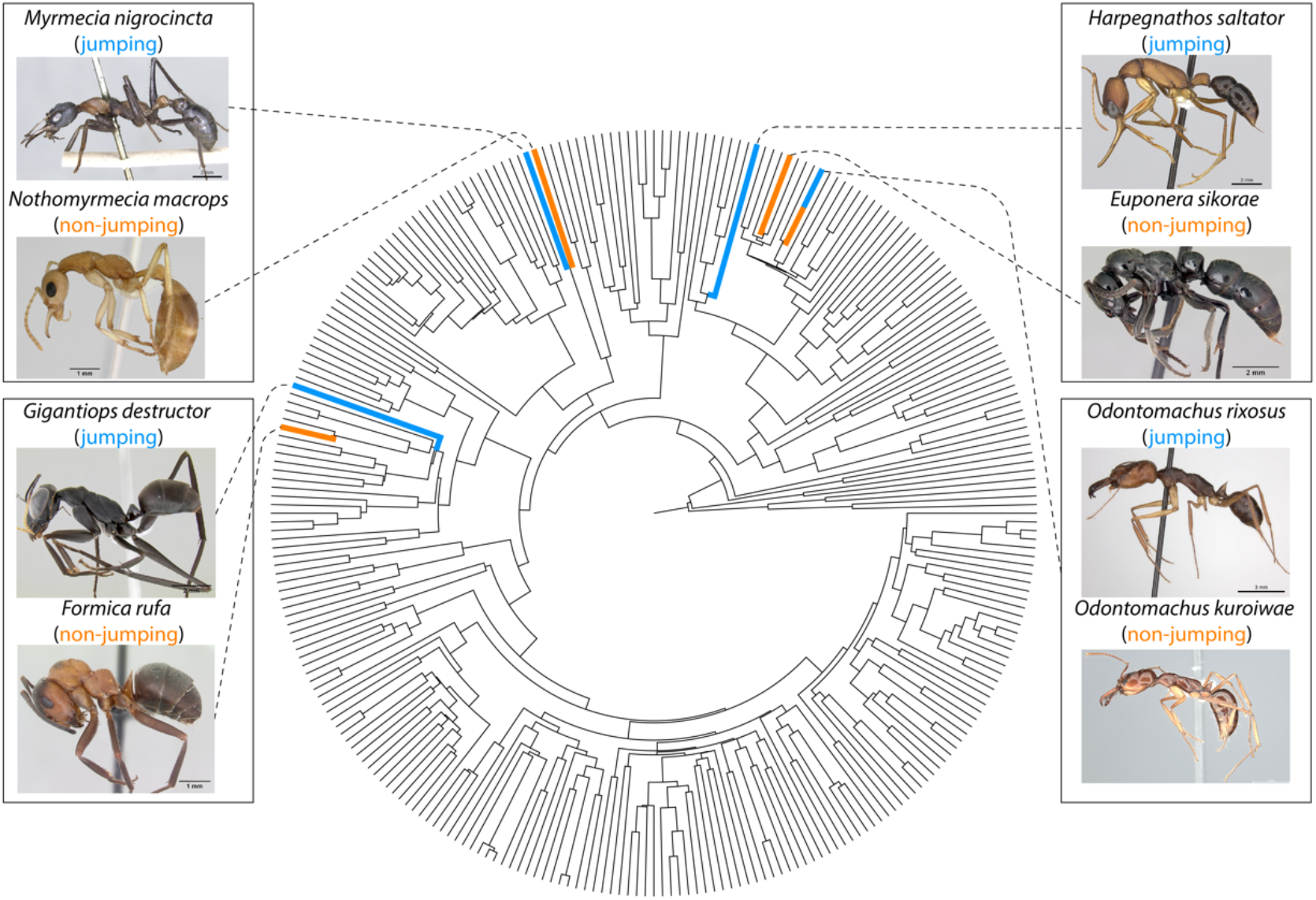
Phylogenetic relationship of the ant species in this study. The phylogenetic tree is from Economo et al. (2018) (the tree was trimmed to include one species per genus). Ant images from Antweb (antweb.org). (*E. sikorae*: CASENT0497202, *F. rufa*: CASENT0173862, *G. destructor*: CASENT0106169, *H. saltator*: CASENT0260424, *M. nigrocincta*: CASENT0902805, *N. macrops*: CASENT0172003, *O. rixosus*: CASENT0217544, *O. kuroiwae*: CASENT0741360).

Most studies on prosalience in ants have focused on the kinematics and energetics of the movements (Ali et al., 1992; Baroni Urbani et al., 1994; Tautz et al., 1994; Ye et al., 2020). In contrast, there has been little fundamental research on jumping abilities in ants from a functional morphology perspective. Although we know how fast these ants jump, we do not yet understand what the adaptations to this function are, and how they differ from non-jumping relatives. Studying the gross anatomy will help us identify the groups of muscles that are essential to facilitate jumps and how are they modified.

All prosalient ants rely on middle and hind legs to jump (Tautz et al., 1994). According to Burrows (2011), there are at least two reasons for using four legs as opposed to a two-legged jump: First, more legs presumably result in a larger net ground reaction force, and so in improved jump performance. Second, it may help to control rotation of the body, so that most muscle work flows into kinetic energy of the Center of Mass (CoM) instead. It has been suggested that the front leg is of minor importance in jumping, and instead acts as a support for maintaining static equilibrium as in case of walking or running (Full & Tu, 1991; Zollikofer, 1994). Although all forward jumping ants universally use the middle and hind legs to jump, the engagement of different body parts during the jump may vary. For example, *Harpegnathos saltator* first uses the hind legs to move the body forward, and only later engages the middle legs to provide a final push (Tautz et al., 1994). However, Baroni Urbani et al. (1994) have shown that the muscle activities in the ipsilateral (same side) middle and hindlegs are synchronous, and thus have concluded that both legs extend in unison. *Gigantiops destructor*, in addition to using both middle and hind legs for propulsion, show a conspicuous and consistent movement of the metasoma (Tautz et al., 1994; Ye et al., 2020), and although the functional significance of this movement is unclear, its consistency suggests it is important. Lastly, *Myrmecia nigrocincta* uses both the middle and hind legs simultaneously to propel the jump (Tautz et al., 1994). As the forward jumping ability of *Odontomachus rixosus* was only recently discovered (Sorger, 2015), the temporal engagement of legs during its jump has not yet been studied.

The involvement of different body parts during jumping may be related to different strategies that would minimize net torque. The position of their Center of Mass (CoM) in this case is important, since the center of rotation is often located in the CoM. This is supported by the observation that the CoM location differs between jumping ant species (Tautz et al., 1994). The impulsive forces generated by the feet in ground contact may be converted into rotational and/or translational kinetic energy (Goode & Sutton, 2023). Rotational kinetic energy may be a particular problem for small animals, as their mass moment of inertia is relatively smaller. The split in rotational versus translational kinetic energy will be determined by the net torque. Zero rotation will only result if the net torque is zero. In *H. saltator,* the CoM is located in front of the middle leg. The use of the middle legs to give the final propulsion could minimize the net torque. The CoM in *G. destructor* is located posterior to the legs, at the insertion of the petiole to the mesosoma. The rotation of the metasoma could be to shift the CoM dynamically to minimize net torques generated by the legs. A similar behavior is observed in juvenile wingless mantises, which rotate their abdomen during their jump to adjust the mass moment of inertia (Burrows et al., 2015). In *M. nigrocincta,* the CoM is located between the middle and hind legs, and perhaps the use of both legs would reduce the total net torque. As such, the differences in the jumping techniques in these ants could be due to the differences in the position of the CoM (Tautz et al., 1994).

There are two size-specific mechanisms used by jumping animals: one powered by direct muscle contraction and one relying on spring-actuated jumps (Sutton et al., 2019). In muscle-actuated jumps, jumping performance is constrained by the physiological properties of the muscle, such as its work density and intrinsic shortening speed. Thus, a closer look at the muscle architecture and volume may elucidate the muscular design for optimal force production.

As animals get smaller in size, the amount of mechanical energy that can be generated is instead limited by the force-velocity properties of the muscle, and it can become beneficial to rely on specialized morphological adaptations that improve performance through rapid recoil of elastic structures (Bobbert, 2013; Sutton et al., 2019). Often, jump enhancement by elastic energy storage involves latch mechanisms: muscles contract slowly to store strain energy in specialised cuticular structures, and this energy is subsequently rapidly released to power the jump (Bennet Clark, 1975; Burrows, 2003; Gronenberg, 1996); although such “power amplification” overcomes force-velocity limitations, it nevertheless ultimately depends on the mechanical work output of muscle. Elastic energy stores in insects are diverse. Click beetles store energy in a specialised structure located in their thorax ventrally between front and middle legs (Bolmin et al., 2021). Locusts use hind femur muscles to load strain energy into the semi- lunar process (Bennet Clark, 1975). Another energy storage site is a locking mechanism found in the femoro-tibial joint (Földvári et al., 2019). So far, it is unknown whether the evolution of forward jumping in ants involves elastic energy storage mechanisms.

## MATERIALS AND METHODS

### Material

We selected one worker specimen preserved in ethanol (70–99%) from each genus for which jumping behavior has been previously documented, to carry out detailed computed tomography (CT) scanning and 3D morphometry: *Harpegnathos saltator* Jerdon, 1851 (unique specimen identifier: CASENT0764679)*, Gigantiops destructor* Fabricius, 1804 (CASENT0709414)*, Odontomachus rixosus* Smith, F., 1957 (CASENT9741319) and *Myrmecia nigrocincta* Smith, F., 1858 (CASENT0741302). As a control, we imaged a small set of non-jumping ant species: *Euponera sikorae* Forel, 1891 (CASENT0709898)*, Formica rufa* Linnaeus, 1761 (CASENT0741323), *Odontomachus kuroiwae* Matsumura, 1912 (CASENT0741313) and *Nothomyrmecia macrops* Clark, 1934 (CASENT0795539). An effort was made to select comparison species as closely related as possible to the jumping species considering availability of preserved specimens. All specimens were stained in iodine solution 3–7 days prior to the scanning date. The length of the leg segments was measured three times within the same specimen (in the right and/or left legs). For the specimens for which no information on the body mass was available, the whole body was scanned, and body volume was used as proxy for body mass via assuming a uniform density of 1040 *kg*/*m*^3^ : *Euponera sikorae* (CASENT0741359); *Formica rufa* (ANTSCAN, CASENT0709272); *Gigantiops destructor* (CASENT0744574); *Nothomyrmecia macrops* (CASENT0741364).

### Micro-CT scanning and 3D-reconstruction

Micro-CT scans were generated with a Zeiss Xradia 510 Versa 3D X-ray microscope operated with the Zeiss Scout-and-Scan Control System software (version 14.0.14829.38124) at the Okinawa Institute of Science and Technology Graduate University, Japan. Scans were conducted with a 40 kV (75 μA) / 3 W beam strength under a 4x magnification. Voxel size and exposure time depended on specimen size (Suppl. Table S1). As the mesosoma of ants exceeds the field-of-view of the camera at high magnification, vertical stitching of serial scans was used. 3D reconstructions of the resulting scan projection data were done with the Zeiss Scout-and-Scan Control System Reconstructor (version 14.0.14829.38124) and saved in txm file format. Postprocessing of txm raw data was done with Amira 2019.2 (Visage Imaging GmbH, Berlin, Germany) to segment individual structures into discrete tissue volumes. The segmented voxels were then exported with the plugin script “multiExport” (Engelkes et al., 2018) in Amira 2019.2 as 2D TIFF image stacks. VG-Studio 3.4 (Volume Graphics GmbH, Heidelberg, Germany) was used to create volume renders from the TIFF image series. Muscle architecture was reconstructed with Amira 2019.2 XTracing extension, following the workflow presented in Katzke, Puchenkov, Stark, & Economo (2022). The accuracy of the tracing algorithm in Katzke et al. (2022) was 92% for fiber length estimation and 100% for the pennation angle estimation. Muscle identity and nomenclature follows Aibekova et al. 2022. Muscles most relevant in the movement of the legs were segmented (Ipcm2, Iscm4, I-, II-, IIIscm1, II-, IIIscm2, I-, II-, IIIscm3, Ipcm8, II-, IIIscm6, Ipcm4, II-, IIIpcm3_4, I-, II-, IIIctm1, I-, II-, IIIctm2, I-, II-, IIIctm3); in addition, large muscles including the indirect muscle of the head (Idvm5), the levator (IA1), and one of the rotators (IA2) of abdomen were segmented for control. We want to mention one caveat related to preparation technique. It is possible there may be different degrees of muscle shrinkage due to the preservation in high ethanol concentrations (70–99%) This could in principle cause spurious differences across species, as the level of contraction may vary among species, however we have no evidence this was an issue in this case, and it is unlikely to affect the broad differences identified in this study.

### Data analysis

Our study design was based on comparing related pairs of jumping and non-jumping species, which should account for phylogenetic signal. However, due to the low throughput of recovering detailed scan and segmentation data for each specimen, and the low number of known independent evolutions of forward jumping (4), we could not perform formal statistical comparative analyses with large numbers of jumping and non-jumping species. Thus, although we quantify morphological differences, our study was mainly performed by comparing values directly, a common limitation of studies of phenotypes without large numbers of independent evolutions. Given the low sample sizes, this exploratory approach can characterize broad and consistent differences but not subtle changes that require large sample sizes. Measurement error and intraspecific variation were assessed however, to ensure our characterization of interspecific differences are not obscured by other sources of variation. For this, we scanned and segmented four *Myrmecia croslandi* specimens from two collection events: *M. croslandi* rep 0 (CASENT0741321), *M. croslandi* rep 1 (CASENT0741324), *M. croslandi* rep 2 (CASENT0741305), *M. croslandi* rep 3 (CASENT0741308). Rep 0 and rep 1 are from an old collection stored in 90% ethanol, Rep 2 and 3 are from the recent collection, stored in 70% ethanol.

In addition to comparing values of morphological parameters (e.g., volumes, fiber lengths) directly, Principal Component Analysis (PCA) was conducted on centered and scaled absolute muscle and thorax volumes to quantify variation in multidimensional space.

### Scaling

In order to compare the morphology across ants of different body sizes, the muscle volume, Physiological Cross-Sectional Area (PCSA), and fiber length were normalized: 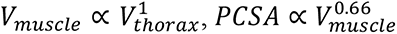, and 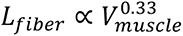. In the absence of detailed information on the force-length properties of the involved muscles (Püffel et al., 2023), we define PCSA as 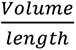. Information on the body mass was lacking for some ants. The data on the volumes of different body parts (head, thorax, petiole, and gaster) of ants, that were calculated from the linear measurements from a single representative species from 231 genera, were available from Anderson, Rivera, & Suarez (2020). We used these data to confirm that thorax volume scales isometrically with total body volume (slope of 1.021 (95% CI: 1.003 - 1.039), Suppl. Fig. S1). Thus, we used the volume of the thorax as a proxy for body mass.

### Data availability

Supplemental Table S1 contains all the specimen data used in this study. Each specimen can be traced by a unique specimen identifier included in the preservation vial. The original µCT scans are available in DICOM format as well as the raw data on the muscle volume, PCSA, pennation angle and fiber length are available at the Dryad Digital Repository. In addition, we have provided freely accessible 3D models of the mesosoma segmentations of all species studied on Sketchfab.

## RESULTS

### Difference in relative volume of muscles involved in jumping

To understand the mechanisms underlying the jumping ability of ants, we compared the normalized muscle size and architecture of several key muscle groups between four jumping ant genera and four closely related non-jumping ant genera. All four jumping ant genera showed changes in size-specific muscle size and architecture of several key muscle groups. Within the meso- and metathorax of jumping species, the trochanter depressor muscles (IIscm6 – *M. mesofurca-trochanteralis*; IIIscm6 – *M. metafurca-trochanteralis*) occupy a large portion of the cavity space (Fig. 2 & Suppl. Fig. S2). In *G. destructor* the relative volume of the trochanter depressor muscle is 5 times larger in the middle legs and 3.5 times in the hind legs compared to *F. rufa*; in *H. saltator* it is 8 times larger in the middle and 14 times larger in the hind legs compared to *E. sikorae*; in *M. nigrocincta* it is 4.5 and 6.5 times larger in the middle and hind legs compared to *N. macrops*; and in *O. rixosus* it is 2.5 and 2 times larger in the middle and hind legs compared to *O. kuroiwae*. The trochanter depressor muscle originates at the furcal arms, the anterodorsal pleural region, and the notum of the meso- and metathorax; it inserts on the trochanter via a long tendon. The homologous muscle in the front leg (Ipcm8) is of similar relative volume in *H. saltator* and *O. rixosus* compared to their non-jumping counterpart, and in *G. destructor* and *M. nigrocincta* it is two times larger compared to their non-jumping counterpart (Suppl. Fig. S3).

**Figure 2.**
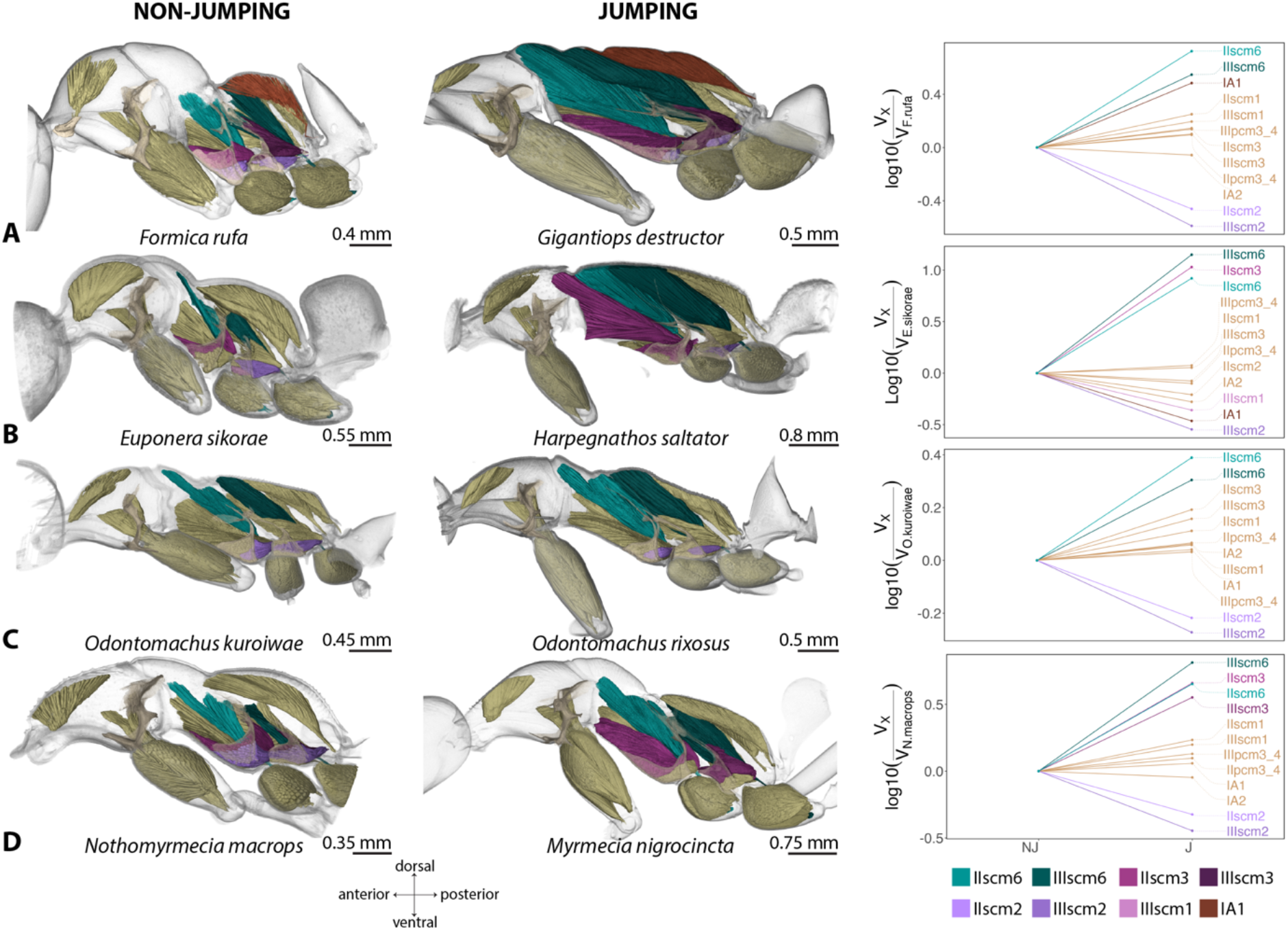
Comparison of the muscle volumes of the leg (coxal and trochanter) muscles. **A**. *F. rufa* and *G. destructor* pair; **B**. *E. sikorae* and *H. saltator* pair; **C**. *O. kuroiwae* and *O. rixosus* pair; and **D**. *N. macrops* and *M. nigrocincta* pair. The line graph on the right side shows the change in the relative volume of the muscles in jumping ant. Muscles which changed the most are colored. The trochanter depressor (scm6) muscles are enlarged in all jumping ants in both middle and hindlegs, while one of the coxal remotor (scm2) muscles are reduced in middle and/or hindleg. *G. destructor* in addition to using the middle and hind legs synchronously, rotate its metasoma to jump; *H. saltator* first uses its hind legs to move the body forward, then the middle legs to give final propulsion; *M. nigrocincta* uses the middle and hind legs synchronously to jump.

Another muscle that is enlarged in relative volume in jumping ants compared to non- jumping ants is one of the remotors of the coxa (IIscm3 – *M. mesofurca-coxalis medialis*; IIIscm3 – *M. metafurca-coxalis medialis*). In *O. rixosus,* scm3 of both middle and hind legs are around 1.5 times larger than in *O. kuroiwae*. In *H. saltator*, scm3 of the middle legs is 10 times larger than in *E. sikorae*, but in the hind legs it is slightly smaller (0.8 times). In addition, the levator muscle of petiole (IA1) is 3 times larger in *G. destructor* compared to *F. rufa,* but the relative volume of IA1 in other pairs is similar.

IIscm2 (IIscm2 – *M. mesofurca-coxalis posterior*) and IIIscm2 (IIIscm2 – *M. metafurca- coxalis posterior*), which are remotors of the coxa (along with scm3 muscles) are reduced in jumping ants (0.2–0.61 times smaller), compared to the non-jumping pairs. 0.35 and 0.26 times smaller in *G. destructor*; 0.62 and 0.28 times smaller in *H. saltator*; 0.47 and 0.36 times smaller in *M. nigrocincta*, and 0.61 and 0.53 times smaller in the middle and the hind legs of *O. rixosus* compared to their non-jumping counterparts.

PCA on the absolute volumes of the muscle and thorax, has shown the separation of jumping and non-jumping ant species on Principal component 1 (PC1) axis; it explained 62.02% of the variation in the sample and PC 2 explained 26.85% of variation in the sample. The most of the variation on PC1 axis come from three muscles: trochanter depressor muscles of the middle and hind legs (IIscm6 and IIIscm6) and coxal remotor muscle of the middle leg (IIscm3).

### Muscle architecture

#### Physiological Cross-Sectional Area

The trochanter depressor muscles of the front (Ipcm8), the middle (IIscm6), and hind legs (IIIscm6) of jumping ants have larger relative Physiological Cross-Sectional Area (PCSA) values (normalized to 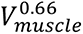 ) compared to non-jumping counterparts (Fig. 3A & Suppl. Table S2), except in the trochanter depressor muscle of the front legs (Ipcm8) of *O. rixosus* and *M. nigrocincta*, where the relative PCSA of was smaller. In *G. destructor* the relative PCSA of the trochanter depressor muscle of the front, middle and hind leg was 1.91, 1.30 and 1.57 times larger, respectively, compared to *F. rufa*. Similarly, in *H. saltator,* the relative PCSA of the trochanter depressor muscle of the front, middle, and hind legs were 1.39, 1.24, and 1.25 times larger compared to *E. sikorae*. In *O. rixosus*, in turn, the relative PCSA of the trochanter depressor muscle the front, middle, and hind legs were 0.74, 2.67, and 2.03 times larger compared non-jumping *O. kuroiwae*. In *M. nigrocincta*, the relative PCSA of the trochanter depressor muscle of the front, middle, and hind legs were 0.94, 1.40, and 1.65 times larger compared to *N. macrops*.

**Figure 3.**
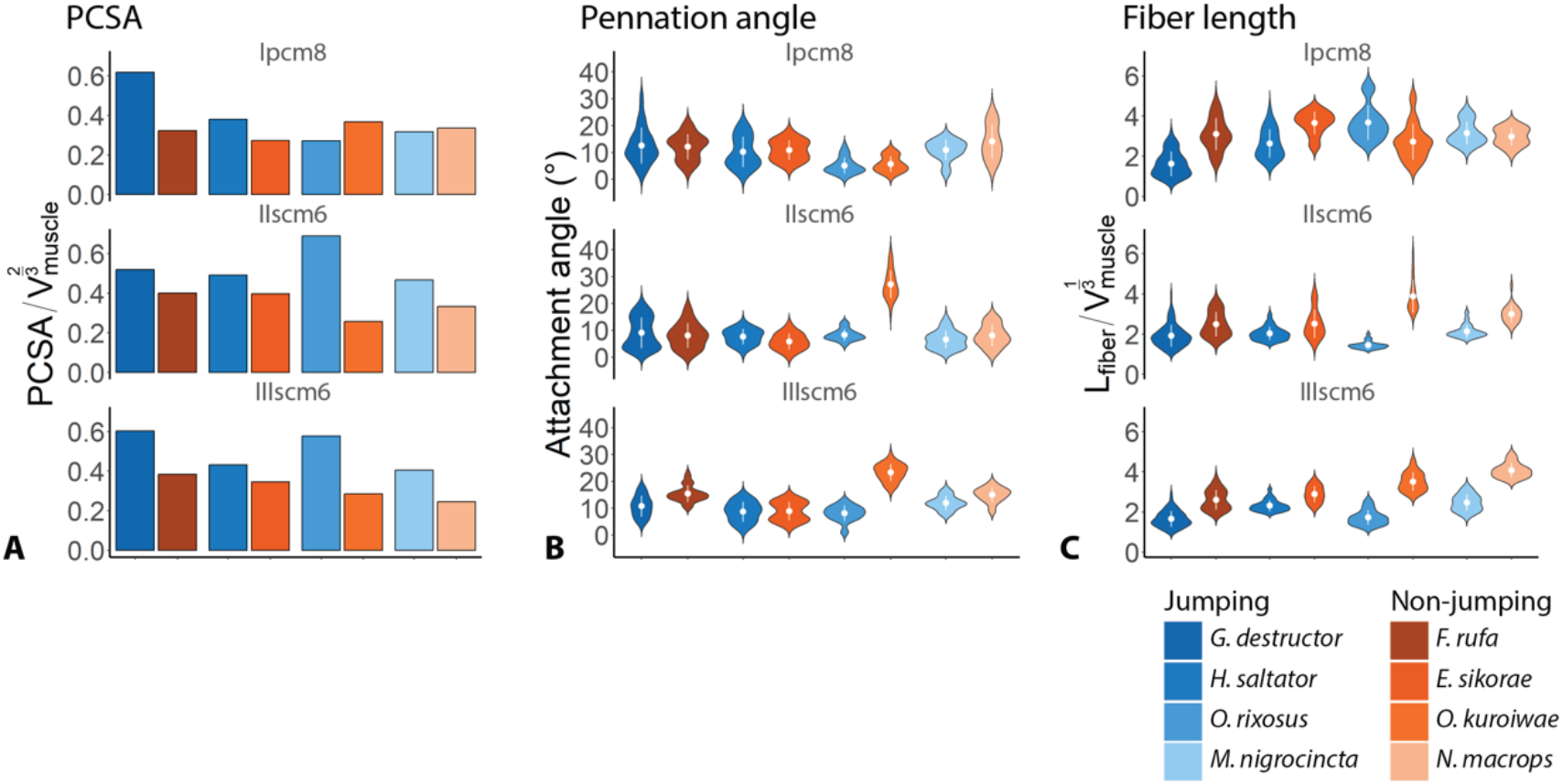
Muscle architecture estimations following fiber tracing of trochanter depressor muscle in the fore (Ipcm8), middle (IIscm6), and hind (IIIscm6) legs. **A**. Effective Physiological Cross-Sectional Area (_EFF_PCSA). To normalize for ant size, we divided the _EFF_PCSA by the approximation of the Surface Area (SA) of the thorax; **B**. Violin plot of the distribution of attachment (pennation) angles, the white dot indicates the mean value, and the white line indicates the s.d.; **C**. Violin plot of the distribution of muscle fiber length, the white dot indicates the mean value, and the white line indicates the s.d.

#### Pennation angle

The mean pennation angle (±s.d., N – number of muscle fibers) in the trochanter depressor muscle in *Gigantiops* and *Formica* pair was similar in the front (13±7, N=77 and 12±5, N=45 accordingly) and middle legs (9±6, N=210 and 8±5, N=31 accordingly). In the hind legs, in turn, it was smaller in *G. destructor* (11±4, N=72), compared to *F. rufa* (15±3, N=39) (Fig. 3B & Suppl. Table S2). Similarly, in *Harpegnathos* and *Euponera* pair, the pennation angle in the front (10±6, N=29 and 11±4, N=17 respectively) and hind legs (9±4, N=91 and 9±3, N=18 respectively) was similar, while in the middle legs it was slightly larger in *H. saltator* (8±3, N=97), compared to *E. sikorae* (6±3, N=19). The mean pennation angle of the trochanter depressor muscle was smaller in all three legs of *M. nigrocincta* (11±4, N=16 in the front, 7±3, N=46 in the middle, and 12±3, N=23 in the hind legs) compared to *N. macrops* (14±6, N=16 in the front, 8±4, N=26 in the middle, and 15±3, N=11 in the hind legs). In *Odontomachus* pairs, the mean pennation angle in the middle and hind legs was larger in non-jumping *O. kuroiwae* (27±5, N=28 in the middle and 23±3, N=29 in the hind legs), compared to jumping *O. rixosus* (8±2, N=17 in the middle and 8±3, N=39 in the hind legs), while in the front legs it was similar (6±3, N=14 and 5±3, N=21 accordingly).

#### Muscle Fiber length

The normalized mean fiber length (±s.d., N – number of muscle fibers) of the trochanter depressor muscle is shorter in jumping ants compared to their non-jumping counterparts in all three legs, except in the front leg of *O. rixosus* and *M. nigrocincta* (Fig. 3C & Suppl. Table S2). In *G. destructor* the relative mean length of fibers was 1.62 (±0.61, N=77), 1.92 (±0.55, N=210) and 1.66 (±0.40, N=72) in front, middle, and hind legs respectively, shorter than those of *F. rufa* (3.1±0.8, N=45, 2.49±0.6, N=31, and 2.6±0.49, N=39). In *H. saltator,* the relative length of muscle fibers 2.63 (±0.71, N=29), 2.03 (±0.34, N=97), and 2.32 (±0.28, N=91) in the front, middle, and hind legs, while in *E. sikorae* it was 3.66 (±0.57, N=17), 2.52 (±0.71, N=19), and 2.89 (±0.4, N=18) accordingly. The relative fiber length of the muscles was longer in the front legs of *M. nigrocincta* (3.15±0.56, N=16), compared to that of *N. macrops* (2.98±0.44, N=16), while in the middle and the hind legs it was longer in *M. nigrocincta* (2.14±0.32, N=46 in the middle and 2.47±0.4, N=23 in the hind legs), compared to *N. macrops* (3±0.44, N=26 and 4.07±0.37, N=11 in the middle and the hind legs, respectively). Likewise, in *Odontomachus* pair, the normalized length of muscle fibers in the front leg is slightly longer in jumping *O. rixosus* (3.67±0.85, N=21) compared to non-jumping *O. kuroiwae* (2.73±0.9, N=14), while in the middle and hind leg it is more than 2 times shorter (1.45±0.19, N=28 and 1.73±0.37, N=29 in *O. rixosus* and 3.88±0.82, N=17 and 3.51±0.46, N=39 in *O. kuroiwae* middle and hind legs accordingly).

### Leg structure

#### The tibial extensor muscle in the hindlegs

The pairwise comparison of the ratio of the intrinsic leg muscles that control the femoro-tibial joints (Fig. 4) shows that in *Gigantiops, Myrmecia,* and *Odontomachus,* the ratio of the extensor muscle to flexor muscle is larger than in their respective non-jumping pairs. Specifically, the ratio in the *G. destructor* and *F. rufa* pair was 1.08 and 0.38, respectively; in *O. rixosus* and *O. kuroiwae* pair 0.87 and 0.55 respectively, and in the *M. nigrocincta* and *N. macrops* pair it was 0.65 and 0.48, respectively. Thus, jumping ants have relatively larger tibia extensor muscles than non-jumping ants, except for the *H. venator* and *E. sikorae* pair. Here, the ratio of the extensor muscle to flexor muscle was 0.44 and 0.75, respectively. However, the tibial extensor muscles (ftm1) in all ants are smaller than the tibial flexor muscles (ftm2), except for *G. destructor,* where the ratio of extensor muscle to flexor muscles is 1.08, slightly larger than one.

**Figure 4.**
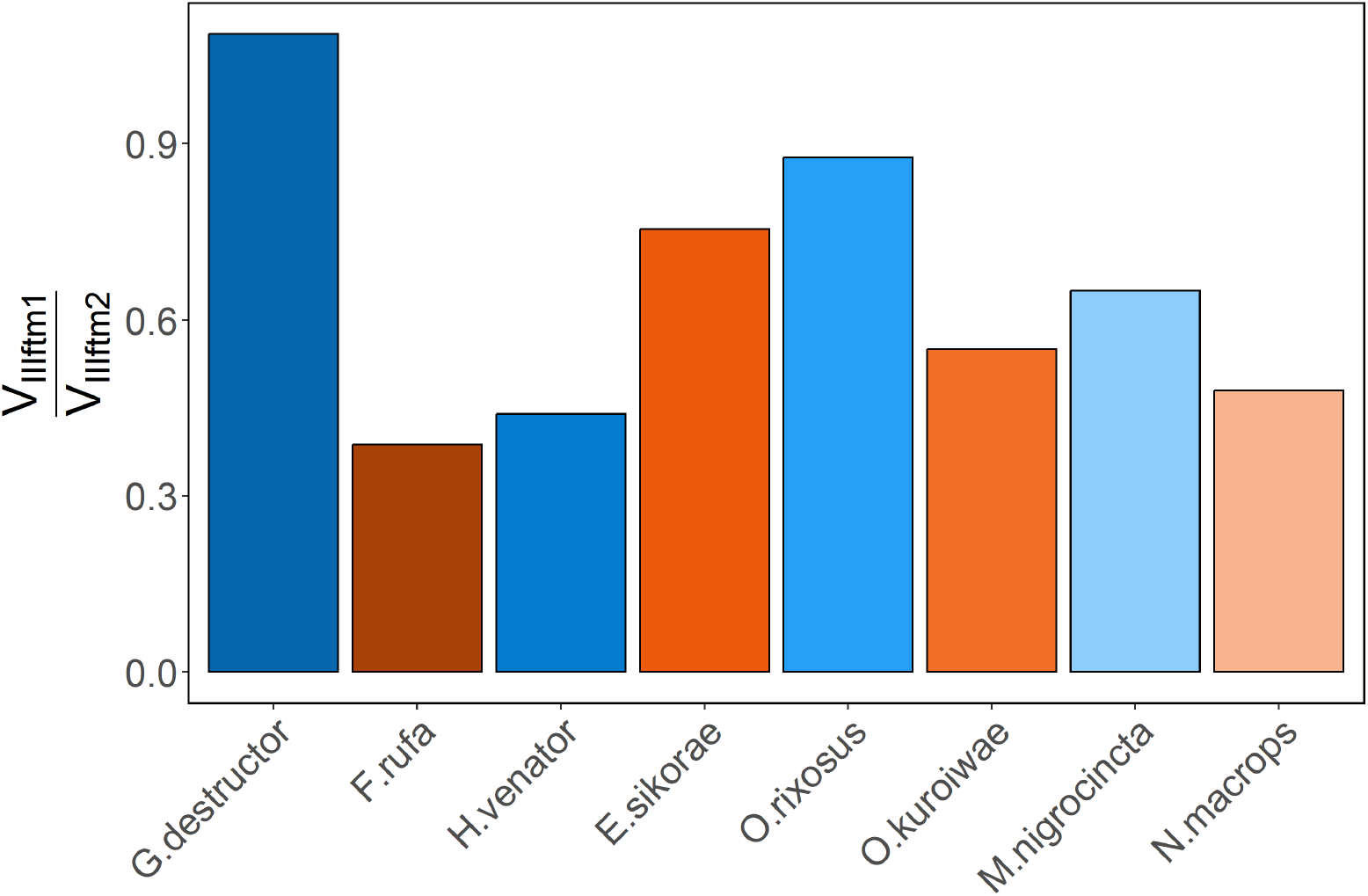
The ratio of the tibial extensor muscle to tibial flexor muscle in the hind legs. The blue bars represent jumping ants and orange bars represent non-jumping ants. The ratio of the extensor muscle to flexor muscle is larger in *G. destructor*, *O. rixosus*, and *M. nigrocincta* than in their non-jumping pairs, expect for *H. venator.* The ratio of the tibial extensor to tibial flexor muscle is smaller than in all ants, except for *G. destructor,* where the volumes of these muscles are almost equivalent.

#### The lengths of legs

In most jumping ants, forelegs are the shortest and hindlegs are the longest (Table 2 & Suppl. Table S3). This is evident in *G. destructor*, where the average length (±s.d.) of the two front legs of was 8.93±0.23 mm, compared to 10.03±0.13 mm and 14.7±0.28 mm for the middle and hind legs, respectively (Suppl. Table S3). The ratio of the leg lengths was thus 1:1.1:1.6 (front:middle:hind) (Table 2). A trend is seen in non-jumping species; for example, in *F. rufa* the average length was 7.08±0.53 mm, 7.24±0.07, 9.03±0.06 mm respectively and the leg length ratio was 1:1:1.3. However, the hind leg length relative to the body length is greater in jumping species, 159% in *G. destructor* and 116% *F. rufa*. Additionally, *G. destructor* have longer femora compared to tibiae in their hind legs, while *F. rufa* have longer tibiae compared to femora.

**Table 2.**
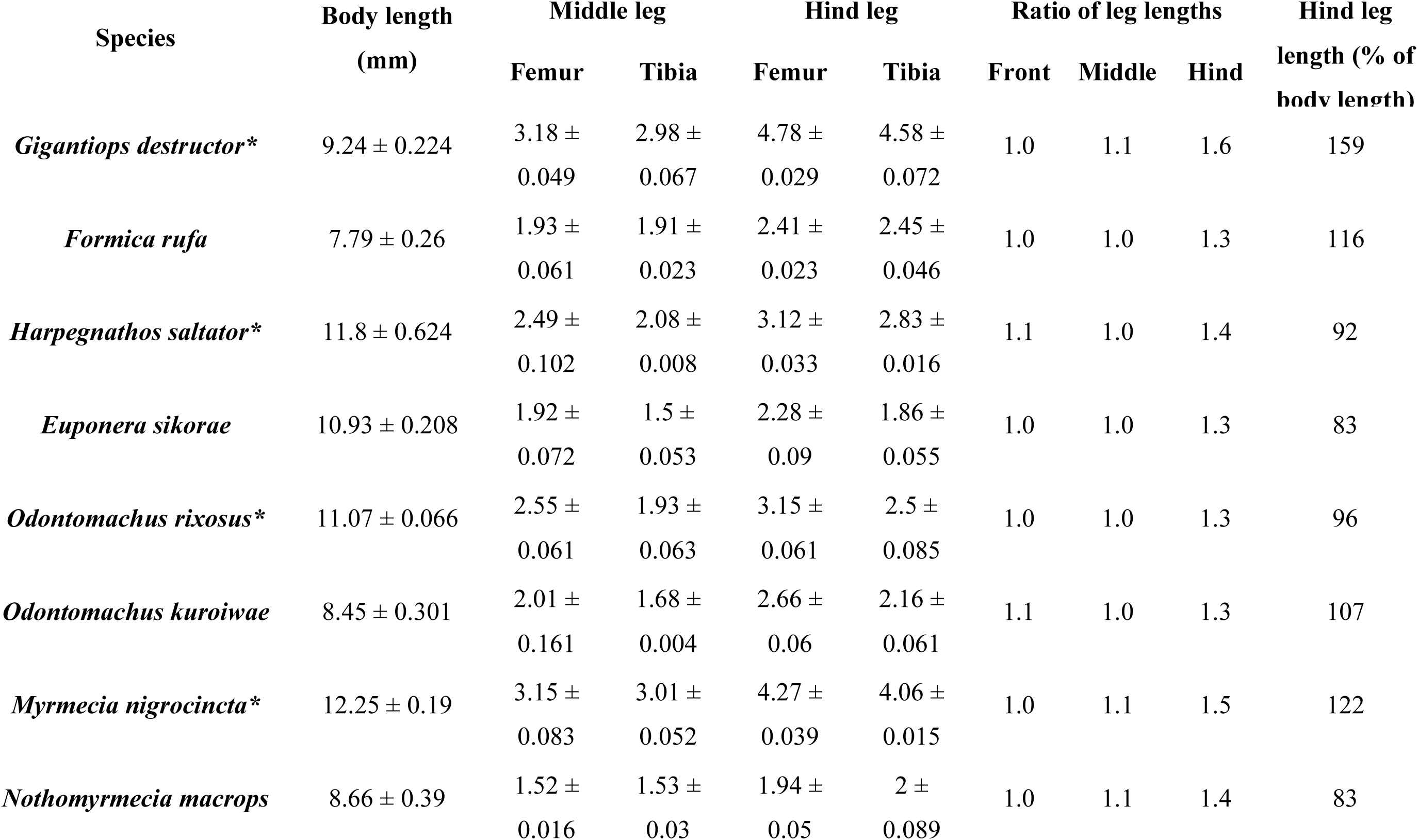
The leg lengths in jumping and non-jumping ants. * indicates jumping ants.

The average length of the front legs of *H. saltator* was 8.46±0.22 mm, the middle legs 7.92±0.04 mm and the hind legs 10.91±0.12 mm (Suppl. Table S3), so the ratio of leg lengths was 1.1:1:1.4. Whereas for *E. sikorae* the respective measurements were 7.19±0.25 mm, 7.13±0.11 mm, and 9.02±0.26 mm, and the ratio was 1:1:1.3. The hind leg length in relation to the body length was 92% in *H. saltator*, while in *E. sikorae* it was 83%.

The average length of the front, middle, and hind legs of *O. rixosus* were 8.63±0.04 mm, 8.22±0.06 mm and 10.62±0.24 mm respectively, as such the ratio of leg lengths was 1:1:1.3. Meanwhile, in *O. kuroiwae* the average leg lengths were 7.46±0.23 mm, 6.9±0.1 mm, 9.01±0.21 mm, resulting in the leg length ratio of 1.1:1:1.3. Additionally, the hind leg length relative to the body length was 96% in *O. rixosus* and 107% in *O. kuroiwae*. Thus, jumping *Odontomachus* have shorter hind legs relative to the body length compared to the non-jumping *Odontomachus*.

The average length of the front, middle, and hind legs of *M. nigrocincta* were 10.15±0.18 mm, 10.94±0.17 mm and 14.93±0.07 mm respectively, resulting in a leg length ratio of 1:1.1:1.5, while in *N. macrops* the average leg lengths were 5.34±0.41 mm, 5.81±0.09 mm, and 7.23±0.08 mm respectively, with a ratio of 1:1.1:1.4. The hind leg length relative to the body length was 122% in *M. nigrocincta*, while it was 83% in *N. macrops*. The non-jumping *N. macrops* has longer tibiae than femora in both middle and hind legs, while in *M. nigrocincta*, the femora are longer than the tibiae (Table 2).

### Intraspecific variation and random error check

From Suppl. Fig. S4, we can see a slight variation in the muscle volume between rep 0–1 and rep 2–3; this variation could be attributed to the ethanol concentration and length of storage time (Leonard et al., 2022; Marquina et al., 2021). However, intraspecific variation is much less compared to interspecific variation. Moreover, a random error check (Suppl. Fig. S5) demonstrates that the error during the segmentation step and computing material statistics is negligible and confirms that the variation in the muscle volumes between species cannot be attributed to measurement error.

## DISCUSSION

*A priori*, the jumping ability in distantly related ant lineages could be associated with different biomechanical and morphological adaptations. However, we found consistent changes whereby trochanter depressor muscles (scm6) in the meso- and metathorax were relatively enlarged across these independent evolutions. Indeed, the first PC (Principal component) axis in the PCA analysis of the absolute muscle volumes separates jumping and non-jumping species rather than grouping them by phylogeny (Fig. 5). Matching the specificities of jumping behavior observed in each lineage, such as stereotypical sequence of movements of different legs or the use of the metasoma, we also uncover a secondary pattern in which other/different groups of muscles most relevant to the behavior/movements are enlarged.

**Figure 5.**
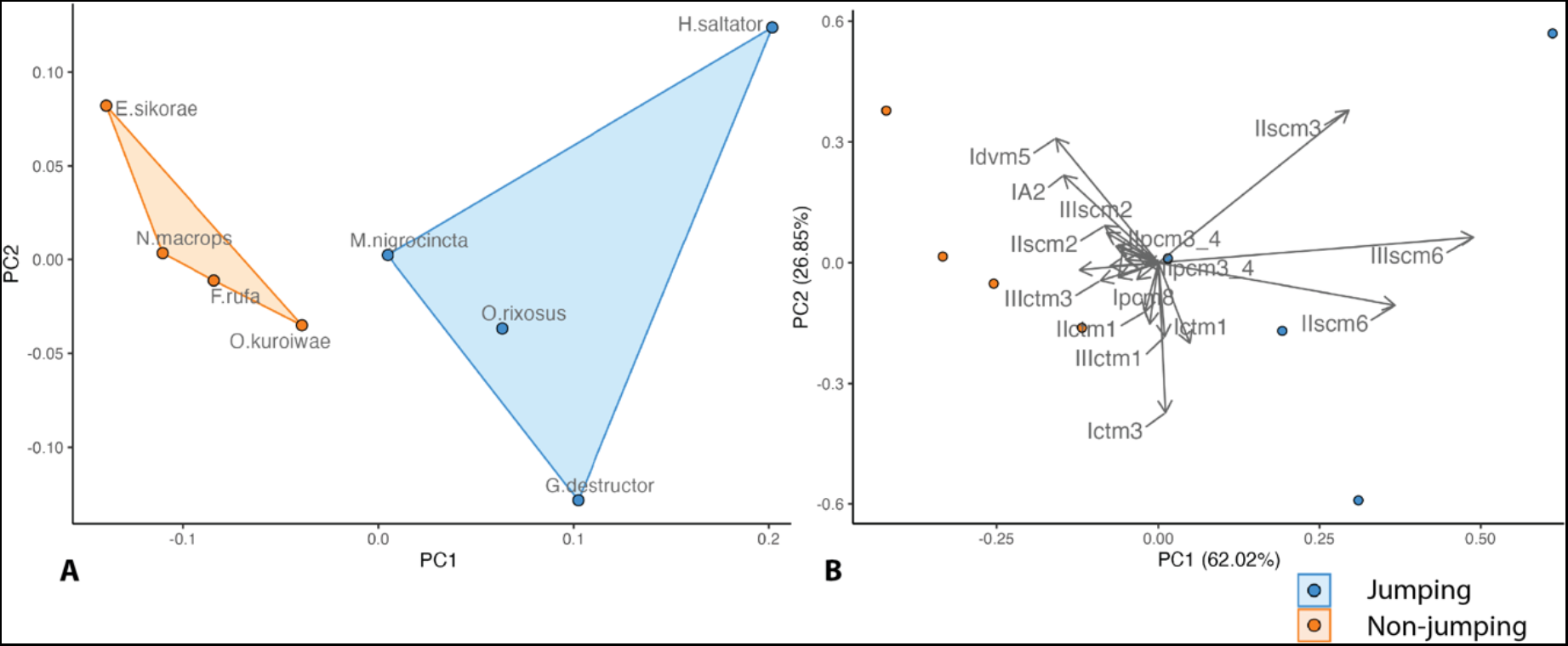
Principal Component Analysis (PCA) of the absolute volumes of muscles and the thorax in ants. **A.** Score plot of the principal components 1 and 2 (PC1 and PC2). **B.** Loadings plot of the PC1 and PC2. The blue dots represent jumping ants and orange represents non- jumping ants. PC1 axis represents 62.02% of the variation, separates jumping and non-jumping ants.

### Changes in relative muscle size

Our results suggest that various muscle groups are modified in jumping ants. The depressor muscles of both mid and hindlegs are greatly enlarged in jumping ants compared to non- jumping ants. Leg depressor muscles are involved in pushing the ant body upward. Conversely, one of the coxal remotor muscles (scm2), small in the first place, are reduced further in jumping ants in both the middle and hind legs, except in the middle leg of *H. saltator*. It is possible that this reduction could be due to the limited space in the rigid mesosoma of ants: enlargement in the volume of one muscle may require the reduction in the volume of another (Fig. 2 & Suppl. Fig. S3). The other coxal remotor muscles (scm3), however, are enlarged in the middle legs of *H. saltator* and in both middle and hind legs of *M. nigrocincta*. In the *Harpegnathos* and *Euponera* pair, the relative volume of the mesocoxal remotor muscles are 10 times larger in the jumping ant. Moreover, these muscles extend to the anterior part of the mesonotum in *H. saltator*, which is unusual for coxal muscles. These changes may be explained by the jumping technique of this genus, which primarily uses the middle legs to jump via a fast “back-rowing” motion (Tautz et al., 1994). However, *M. nigrocincta* uses both legs to jump, which may be the reason why scm3 muscles are enlarged in both middle and hindlegs. The increase in the relative volume of the mesial coxal remotor muscles is not substantial in other jumping ants. Lastly, *G. destructor* which rotates the metasoma while jumping, has relatively larger petiole levator muscles (IA1). These differences in muscle mass suggest that different morphological modifications are related to the observed variation in the jumping technique.

The ratio of the tibial extensor muscle to tibial flexor muscle is less than unity in most ants, which is a characteristic of grasping legs in other insect lineages (Földvári et al., 2019). In *G. destructor,* the ratio is 1.08, so that *G. destructor* has almost equivalent volume of the tibial extensor and flexor muscle, which is a characteristic of walking legs in other insects (Földvári et al., 2019). In *Gigantiops, Myrmecia,* and *Odontomachus* the ratio of the tibial extensor to flexor muscle is in fact larger than that of their non-jumping counterparts. However, the fact that overall, the tibial extensor muscle is smaller than the flexor muscle suggests that ants may not rely as much on the tibial extensor muscle to jump.

If we compare the relative muscle volumes across genera, *Myrmecia* has an enlarged volume compared to the non-jumping sister lineage *Nothomyrmecia*. However, within *Myrmecia,* we see a surprising pattern: non-jumping *Myrmecia* have similar trochanter depressor muscle volumes as jumping *Myrmecia*. It is possible that some *Myrmecia* have cryptic jumping ability. Despite their fame as “jumping jack ants”, studies on the locomotion modes across *Myrmecia* species are fragmentary and contradictory (Brown, 1953; Clark, 1943; Snodgrass, 1942; Wheeler, 1922). In order to better understand this phenomenon, we plan to conduct further research that combines kinematics with analysis of muscle volumes. This will help us to clarify the patterns we have observed and determine whether some Myrmecia species may possess cryptic jumping abilities.

### Changes in muscle architecture

Above we showed that there is an increase in size-specific muscle volume. An increase in muscle volume can be invested in area, in length, or in both. To understand whether jumping ants preferentially invest the additional muscle volume in PCSA or in fiber length, we directly compared the muscle architecture of the trochanter depressor muscles. The relative PCSA of the trochanter depressor muscles of the front (Ipcm8), the middle (IIscm6), and hind legs (IIIscm6) of jumping ants was larger compared to non-jumping counterparts, except in the trochanter depressor muscle of the front legs of *O. rixosus* and *M. nigrocincta*, where the relative PCSA of was smaller. On the other hand, the normalized mean fiber length of the trochanter depressor muscle was shorter in jumping ants compared to their non-jumping counterparts in all three legs, except in the front leg of *O. rixosus* and *M. nigrocincta.* This shows that the jumping ants preferentially invest the increased volume of the trochanter depressor muscle into PCSA, except in the front legs of *O. rixosus* and *M. nigrocincta*.

There is no clear difference in the pennation angle of the trochanter depressor muscles between jumping and non-jumping ants. In *Odontomachus* pairs, the mean pennation angle in the middle and hind legs was larger in *O. kuroiwae* compared to *O. rixosus*, this change in the pennation angle is small: the force “loss” associated with pennation is *cos*(8) = 0.99 in *O. rixosus*, and *cos*(27) = 0.89 in *O. kuroiwae*. Thus, a more than threefold difference in pennation angle only results in about 10% reduction in instantaneous force capacity.

This parallelism among jumping ants with regards to the increase in relative volume of the trochanter depressors scm6 in the mid and hindlegs is particularly relevant due to the anatomical peculiarities of this muscle. Unlike the other trochanter depressor (ctm3) which originates inside the coxa (as do the trochanter levator pair, ctm1–ctm2), trochanter depressor scm6 is an extrinsic muscle which originates in the thorax (Aibekova et al., 2022). This general anatomical arrangement already results in the muscle being longer than any of the other trochanter muscles in non-jumping ants, while in jumping ants it was possible for this muscle to enlarge into proportions that are effective for upward action, occupying a large portion of the otherwise free thoracic cavity. Moreover, being extrinsic to the coxa, the action of muscle scm6 is achieved independently of the promotion and remotion of the leg controlled by the coxal muscles. The result is that jumping ants can effectively swing the mid and hindlegs backwards to push their body forward while at the same time generating the strong upward lift from the extrinsic leg depressor transmitted from the thorax to the trochanter by way of the long scm6 tendon.

### Jump mechanism– elastic recoil or direct muscle contraction?

We have identified consistent differences in the relative volume and architecture of leg muscles across jumping and putatively non-jumping ants. We now assess if observed jump performance can be explained by direct contraction of these muscles, or whether contribution from recoiling elastic elements is required. To this end, we first calculate the total work and average power requirements of observed jumps from published data, 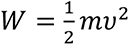, and 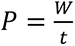, respectively; here, *m* is the body mass, *υ* is the peak take-off velocity, and *t* is the take-off time (see Table 1). We neglect gravitational potential energy in these calculations, as it only makes a marginal contribution in small animals (see Labonte, 2022; Scholz et al., 2006). Next, we estimate the muscle volume required to achieve this work or power output, making use of the observation that the work and average power density – the maximum amount of work and average power a unit muscle mass can generate – are relatively conserved across disparate taxa, *W_ρ_*: ≈ 70 J/*kg* and P*_ρ_*: ≈ 350 W/*kg* (Alexander R McNeill, 2003; Biewener & Patek, 2018; we assume a muscle density of 1040 *kg* /m^3^). In a last step, these volume estimates may then be compared to the measured muscle volume for the trochanter depressor muscles of middle and hind legs. While it is possible that multiple muscles can be involved in powering the jump, our estimation was based on the assumption that the trochanter depressor muscles provide the primary force for upward lift; thus, we did not factor in the power output from the other leg muscles. Comparison of the power-based muscle volume assesses the possibility that jumps could be driven by direct muscle contraction; comparison of the muscle volume estimated via the work requirements, in turn, probes the possibility that the jump may have been meaningfully enhanced by recoil of elastic elements. Previous work has argued that muscle can convert at most a third of its work density into elastic strain energy, provided that the spring constant of the involved spring-like elements takes its optimal value (Sutton et al., 2019); we thus multiply the work estimate by three, to derive a conservative lower bound for the required muscle volume. These simple calculations suggest one clear case (see Table 1): For *M. nigrocincta*, leg muscles have just about enough work capacity to drive a jump via intermediate conversion of muscle work into elastic strain energy, but their power capacity falls short by about a factor four; jumps in this species are thus likely enhanced by elastic recoil. While our estimation accounted for the contribution of the trochanter depressor muscles only, it is worth noting that the observed enlargement of the coxal remotor muscles (scm3) in both middle and hind legs suggest that they might also contribute to the jump of *Myrmecia*. For both *G. destructor and in H. saltator,* in turn, the estimates are less conclusive. In both species, the total muscle volume would suffice to power the jumps directly, or to drive them via elastic recoil. Indeed, Baroni Urbani et al. (1994) assumed that jumps of *H. saltator* are powered solely by muscle, on the basis of the low take-off velocity; but they also did not exclude the possibility of energy storage. We looked for evidence for any of the three locking elements identified by Földvári et al. (2019) in the micro-CT scans of the femoro-tibial joints in all jumping species; none were found in any of the jumping ants (Suppl. Fig. S6). Similarly, bush crickets, which use direct muscle contraction to power their jumps, do not possess a locking mechanism (Földvári et al., 2019). Although this suggests that jumps in *G. destructor* and *H. saltator* may indeed be driven by direct muscle contraction alone, it leaves open the question how and which elastic elements contribute to jump performance in *M. nigrocincta*; perhaps these ants do not rely on latching, but load elastic elements dynamically during the jump, as is the case in larger jumping vertebrates such as frogs (Marsh, 1994).

**Table 1.** The parameters for the calculations of the energy requirements.

|  | *G. destructor* | *H. saltator* | *M. nigrocincta* |
| --- | --- | --- | --- |
| **Body mass in kg** | 0.0000155 | 0.000032<sup>a</sup> | 0.0000266<sup>b</sup> |
| **Take-off speed in m/s** | 0.6<sup>c</sup> | 0.7<sup>c</sup> | 0.96<sup>b</sup> |
| **Take-off time in s** | 0.0275<sup>c</sup> | 0.02<sup>c</sup> | 0.0175<sup>b</sup> |
| **Work** | $0.28 \times 10^{-5}$ | $0.78 \times 10^{-5}$ | $1.23 \times 10^{-5}$ |
| **Power** | $0.10 \times 10^{-3}$ | $0.39 \times 10^{-3}$ | $0.70 \times 10^{-3}$ |
| **Muscle volume in m<sup>3</sup> based on work requirements** | $1.15 \times 10^{-10}$ | $3.23 \times 10^{-10}$ | $5.05 \times 10^{-10}$ |
| **Muscle volume in m<sup>3</sup> based on power requirements** | $2.79 \times 10^{-10}$ | $1.08 \times 10^{-9}$ | $1.92 \times 10^{-9}$ |
| **Measured total muscle volume** | $4.1 \times 10^{-10}$ | $1.29 \times 10^{-9}$ | $4.94 \times 10^{-10}$ |

*Note.* Data are from Baroni Urbani et al. (1994)<sup>a</sup>, Allman et al., unpublished data<sup>b</sup>, and Tautz et al (1994)<sup>c</sup>.

It is remarkable that the increase in relative muscle volume appears to be invested dominantly in PCSA (see also Püffel et al., 2021 for a similar result in leaf-cutter ant mandible closer muscle). The functional advantage of this preferential investment is not obvious, because both the work and the power output of muscle depend only on muscle volume. In *M. nigrocincta*, preferred investment in PCSA may be driven by the structural and mechanical properties of the elastic elements, which may require “matching” of muscle properties to maximize the storage of elastic strain energy (Rosario et al., 2016). In *G. destructor* and *H. saltator*, in turn, the increase in peak net force arising from the increase in PCSA would result in shorter take-off times but may leave peak-speed unaffected (Labonte, 2022), suggesting that it serves to enable rapid escape maneuvers. Future work is required to address these conjectures in more detail.

## CONCLUSION

In this study, we describe morphological adaptations associated with the evolution of forward jumping in ants. We found that all jumping ants have enlarged the relative volume of trochanter depressor muscles in the middle and hind legs, as well as a reduced relative volume of the posterior remotor muscle of the coxa. However, different sets of muscles are enlarged based on which body parts are involved in jumping: medial remotor of the coxa of the middle leg in *Harpegnathos*; medial remotor of the coxa of both middle and hindlegs in *Myrmecia*; levator of the petiole and extensor of the tibia in *Gigantiops*. This indicates that forward jumping ants have undergone parallel evolution in a broad sense, producing jumps through modifications of the leg system without latch mechanisms. However, there is secondary variation in both jumping mechanics and musculature as ants vary in their use of hind and mid-legs, which could be considered an example of many-to-one mapping. Based on direct comparison of the observed vs. possible work and power output during jumps, we suggest that direct muscle contractions suffice to explain jumping performance, in *G. destructor* and *H. saltator*, but elastic recoil is likely important in *M. nigrocincta*. These results help to elucidate morphological aspects of how forward jumping has evolved in ants and shed light on how biomechanical systems and behaviors co-evolve to generate diversity.

## ACKNOWLEDGEMENTS

We are deeply grateful to Christian Peeters for bringing the authors from different continents of this study together. We thank the Okinawa Institute of Science and Technology Graduate University (OIST) Imaging Section for providing access to the Zeiss Xradia micro-CT scanner and Shinya Komoto for general support. This work was supported by several Japan Society for the Promotion of Science (JSPS) grants-in-aid KAKENHI grants [No. 21J20268 to LA; No. 17K15180 to EPE; No. 18K14768 & 21K06326 to FHG], and a grant from the Japan Ministry of the Environment (Environment Research and Technology Development Fund no. 4-1904 to EPE) and an Australian Research Council’s Discovery Project grant (DP220102836 to AN).

## SUPPLEMENTAL MATERIAL

**Supplemental Figure S1.**
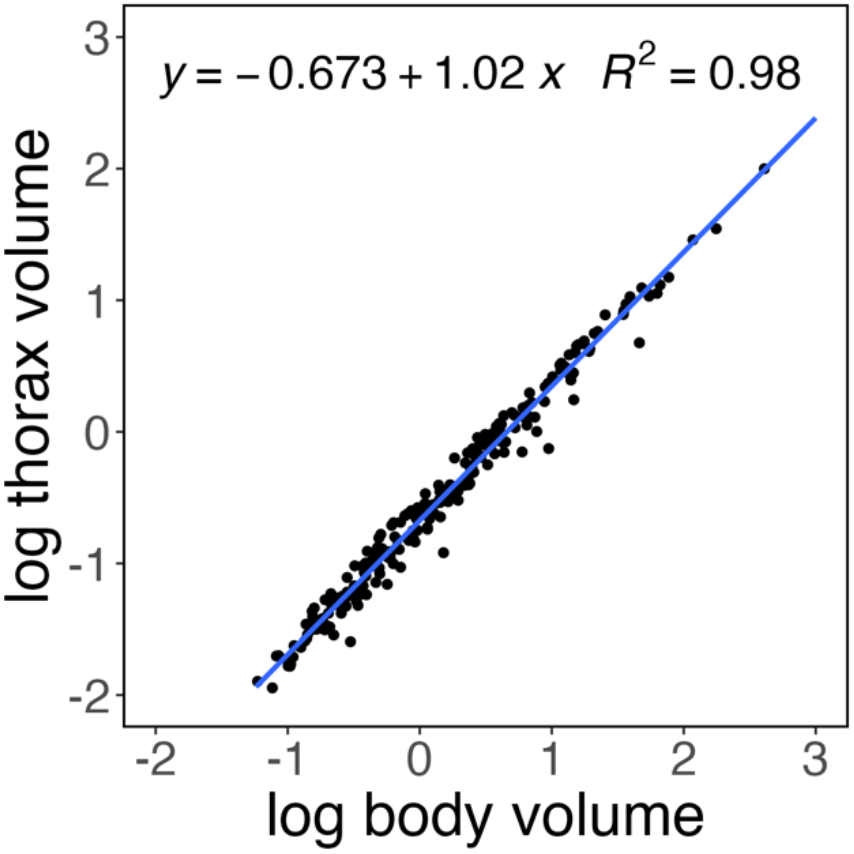
Allometric scaling relationship between the thorax volume and the body volume in ant workers. The thorax volume scaled isometrically with the body volume.

**Supplemental Figure S2.**
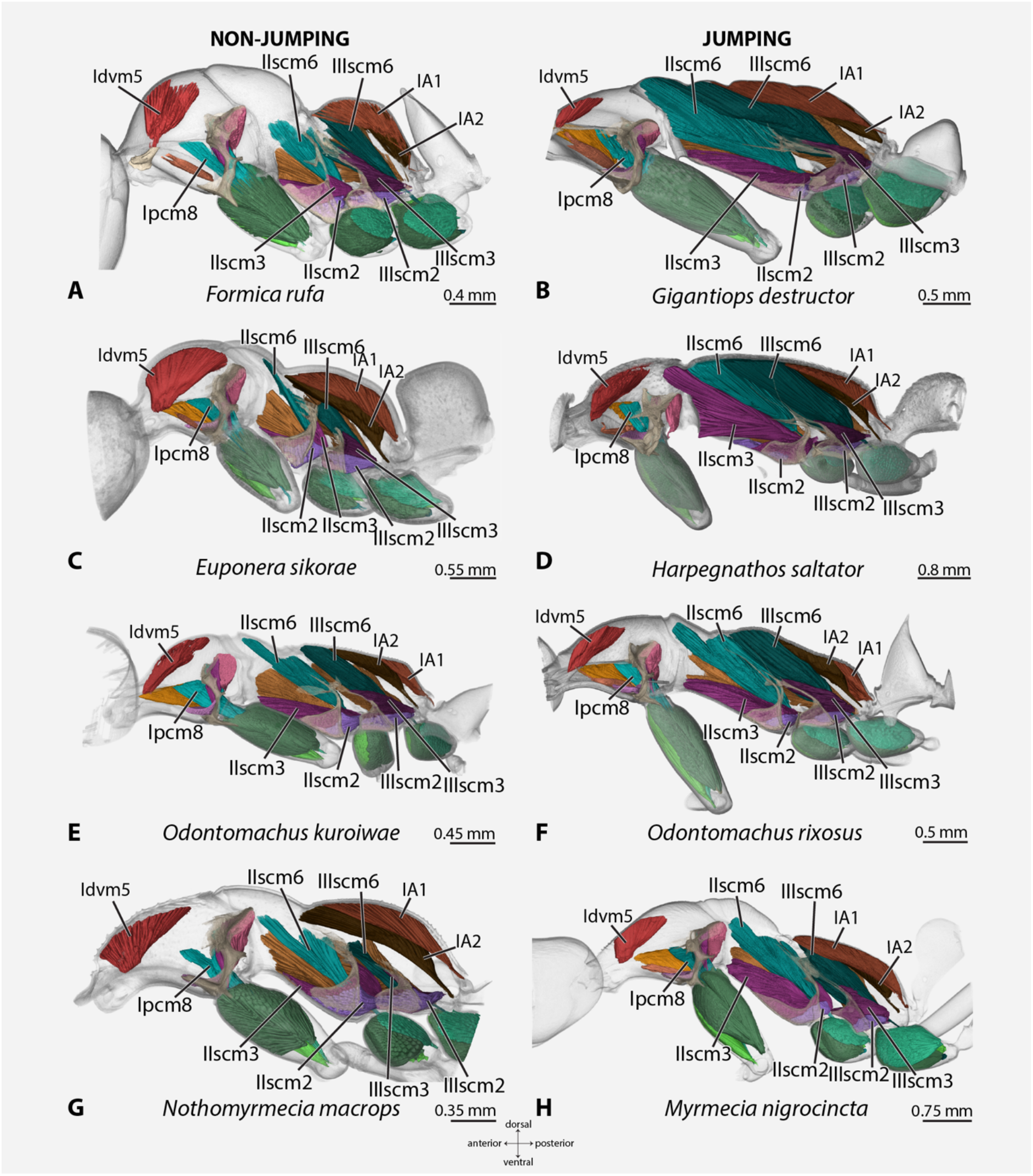
The 3D reconstruction of the mesosoma of ants in Figure 1.

**Supplemental Figure S3.**
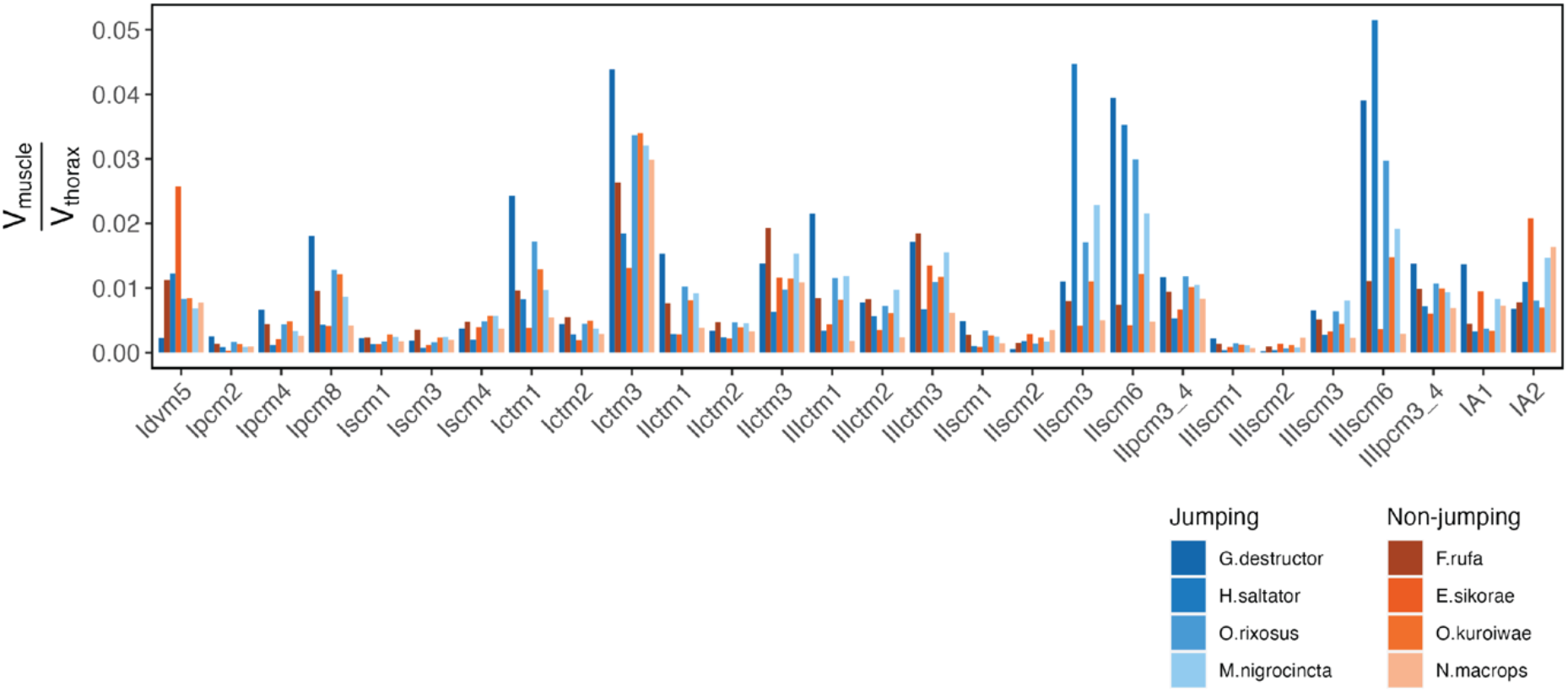
The relative muscle volume comparison of all leg muscles in front, middle, and hind legs, as well as abdominal muscles and a head muscle.

**Supplemental Figure 4.**
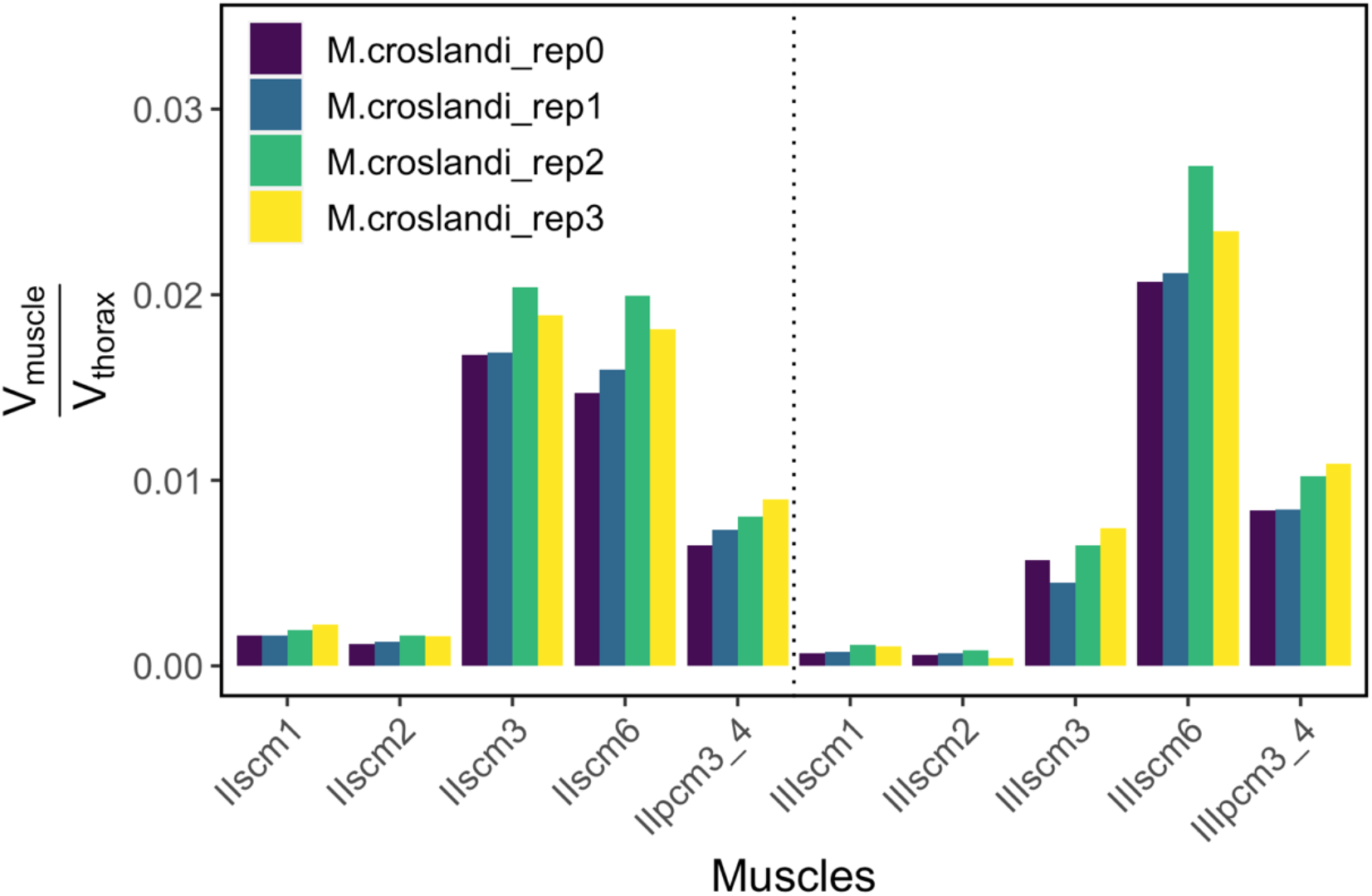
Repetitions of muscle segmentation and muscle volume comparison for *Myrmecia croslandi*. The specimens were picked from different collection events and different alcohol concentration (70% and 90%).

**Supplemental Figure S5.**
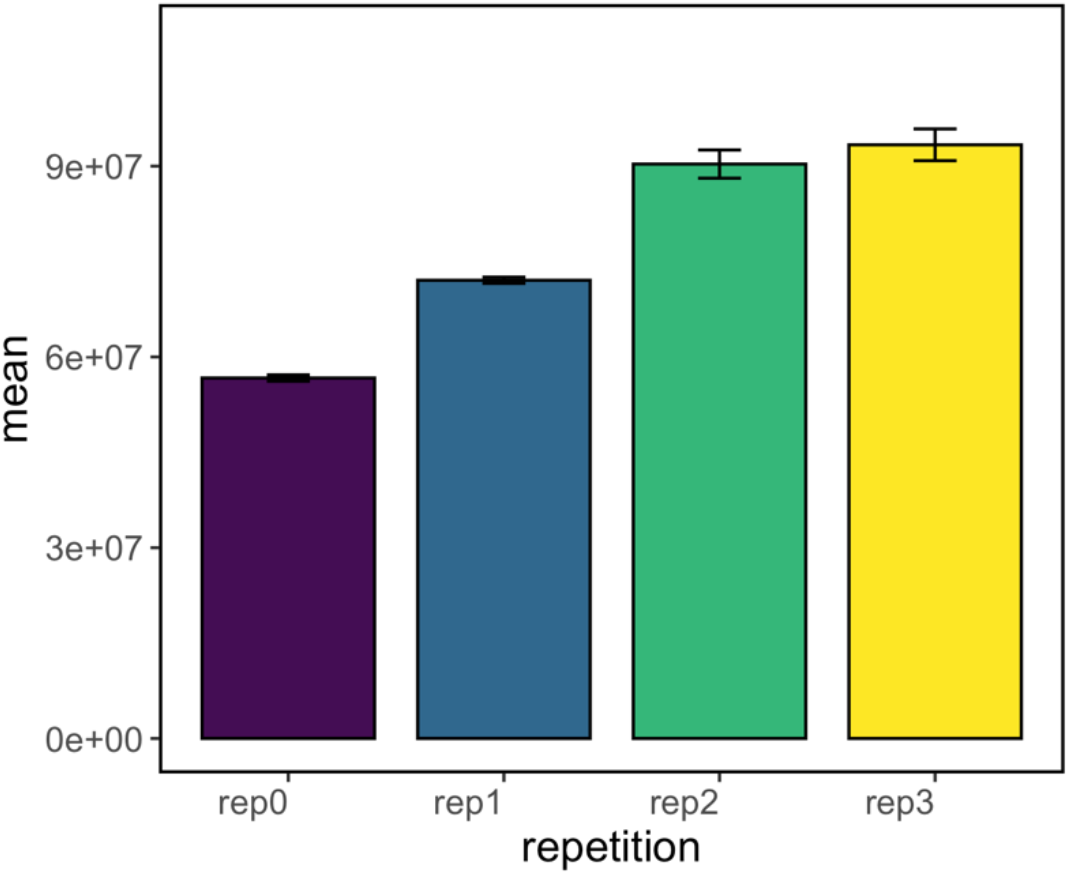
Random error check for the muscle segmentation. IIscm6 muscle was segmented 5 times in each of the repetitions. The error bar indicates the SD.

**Supplemental Figure S6.**
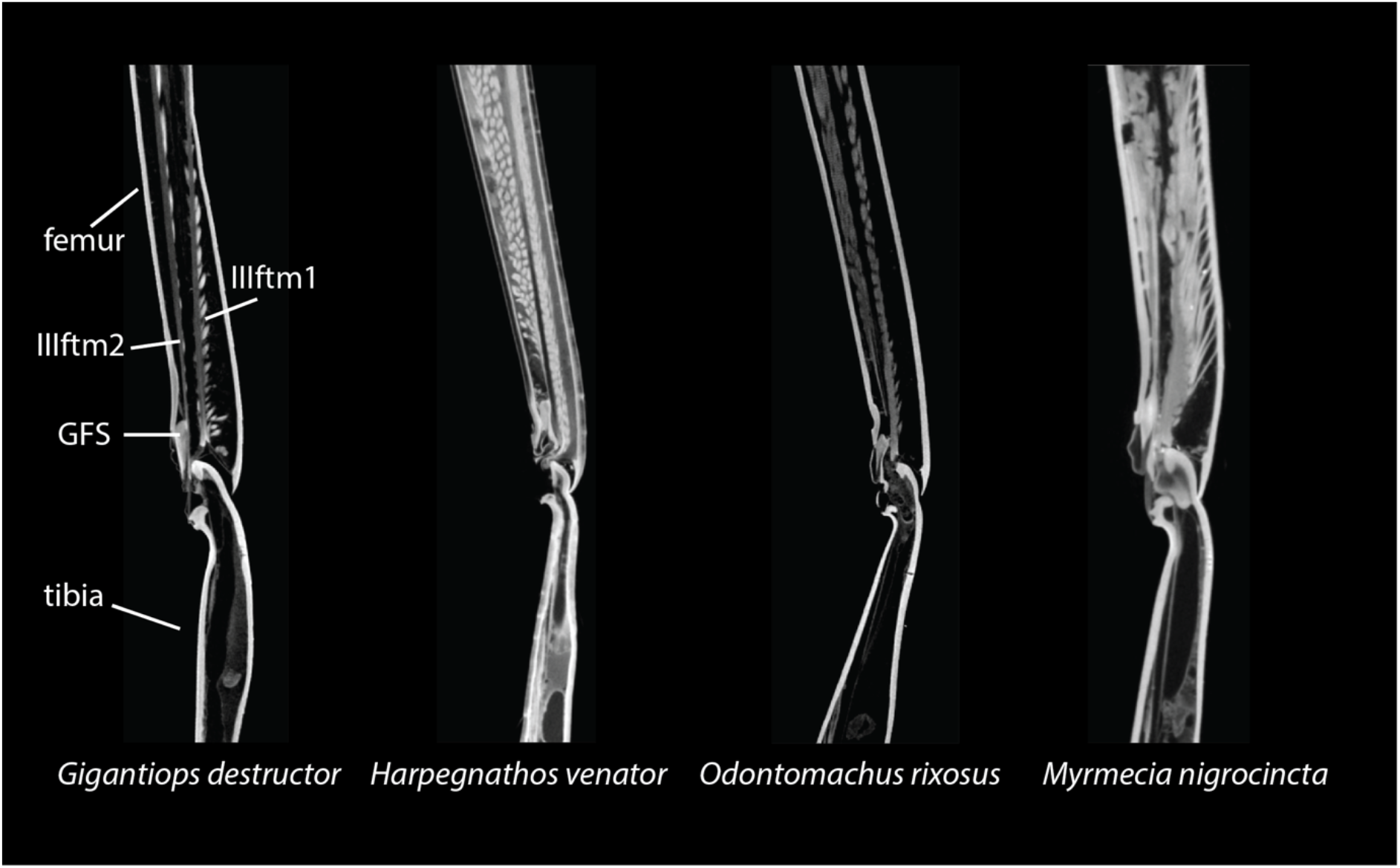
The CT scan image of the femoro-tibial joint in mid-sagittal plane.

**Supplemental Table S1**. The metadata of the specimen and scan settings.

**Supplemental Table S2.** Muscle architecture of trochanter depressor muscles of the front, middle and hind legs.

**Supplemental Table S3**. Leg lengths measurements of the front, middle and hind legs.

